# Distinct lung-homing receptor expression and activation profiles on NK cell and T cell subsets in COVID-19 and influenza

**DOI:** 10.1101/2021.01.13.426553

**Authors:** Demi Brownlie, Inga Rødahl, Renata Varnaite, Hilmir Asgeirsson, Hedvig Glans, Sara Falck-Jones, Sindhu Vangeti, Marcus Buggert, Hans-Gustaf Ljunggren, Jakob Michaëlsson, Sara Gredmark-Russ, Anna Smed-Sörensen, Nicole Marquardt

## Abstract

Respiratory viral infections with SARS-CoV-2 or influenza viruses commonly induce a strong infiltration of immune cells into the lung, with potential detrimental effects on the integrity of the lung tissue. Despite comprising the largest fractions of circulating lymphocytes in the lung, little is known about how blood natural killer (NK) cells and T cell subsets are equipped for lung-homing in COVID-19 and influenza. Using 28-colour flow cytometry and re-analysis of published RNA-seq datasets, we provide a detailed comparative analysis of NK cells and T cells in peripheral blood from moderately sick COVID-19 and influenza patients, focusing on the expression of chemokine receptors known to be involved in leukocyte recruitment to the lung. The results reveal a predominant role for CXCR3, CXCR6, and CCR5 in COVID-19 and influenza patients, mirrored by scRNA-seq signatures in peripheral blood and bronchoalveolar lavage from publicly available datasets. NK cells and T cells expressing lung-homing receptors displayed stronger phenotypic signs of activation as compared to cells lacking lung-homing receptors, and activation was overall stronger in influenza as compared to COVID-19. Together, our results indicate migration of functionally competent CXCR3^+^, CXCR6^+^, and/or CCR5^+^ NK cells and T cells to the lungs in moderate COVID-19 and influenza patients, identifying potential common targets for future therapeutic interventions in respiratory viral infections.

**Author summary:** The composition of in particular CXCR3^+^ and/or CXCR6^+^ NK cells and T cells is altered in peripheral blood upon infection with SARS-CoV-2 or influenza virus in patients with moderate disease. Lung-homing receptor-expression is biased towards phenotypically activated NK cells and T cells, suggesting a functional role for these cells co-expressing in particular CXCR3 and/or CXCR6 upon homing towards the lung.

## Introduction

The ongoing pandemic of coronavirus disease 19 (COVID-19), caused by the novel severe acute respiratory syndrome coronavirus 2 (SARS-CoV-2), highlights the need for a better understanding of respiratory viral infections which have the potential to cause recurrent epidemics or pandemics. In addition to SARS-CoV-2, this also includes influenza virus, respiratory syncytial virus (RSV), SARS-CoV, and the Middle East Respiratory Syndrome (MERS)-CoV. Future disease outbreaks with novel viruses affecting the airways are to be expected and prepared for.

During acute infections with respiratory viruses such as with SARS-CoV-2 or influenza, specific chemokines mediating leukocyte recruitment are increased in the lung and bronchoalveolar lavage (BAL) fluid. These chemokines include CCL2, CCL3, CCL20, CXCL1, CXCL3, CXCL10, and IL8, attracting cells expressing chemokine receptors such as CCR2, CCR5, CXCR3, and CXCR6 (1-3). In patients suffering from severe COVID-19, recent reports suggest exacerbated lung tissue damage and epithelial cell death resulting from hyperactivated immune cells, such as inflammatory macrophages (1), natural killer (NK) cells (4) and/or T cells (1,5). While moderate disease in COVID-19 and in influenza patients is per definition not fatal, patients may still require hospitalization and/or experience persistent long-term symptoms such as fatigue, respiratory problems, loss of taste or smell, headache, and diarrhea. Since lung-homing cytotoxic lymphocytes are likely involved in lung pathology during acute infection, a better understanding of their major homing mechanisms will help in developing and improving treatment strategies in COVID-19 and other respiratory viral infections.

In this study, we investigated the composition of NK cell and T cell subsets which are equipped with lung-homing properties in the peripheral blood of patients suffering from moderate COVID-19 or influenza, and in healthy controls. In addition to analyses by 28-colour flow cytometry, we assessed transcript expression in NK cells and T cells using three publicly available single cell (sc)RNA-seq datasets of cells from peripheral blood and bronchoalveolar lavage, respectively (6-8). Our data indicate a universal role for CXCR3-mediated lung-homing of NK cells and T cells in COVID-19 and influenza and an additional role for recruitment via CXCR6 and CCR5.

Together, we provide an extensive characterization of the lung-homing potential of functional NK cells and T cells in homeostasis and acute respiratory viral infections such as COVID-19 and influenza. The present results are of relevance for the understanding of the disease progression and for identifying target molecules to improve future therapeutic treatment strategies.

## Results

### Altered frequencies of chemokine receptor-positive NK cell and T cell subsets in peripheral blood in COVID-19 and influenza patients

Chemokine receptors relevant for lung-homing such as CXCR3, CXCR6, CCR2, and CCR5, could be identified in all detectable NK and T cell subsets in peripheral blood both from healthy donors and COVID-19 patients (Fig. 1a, b; see Fig. S1a for gating strategies). The frequency of chemokine receptor-positive NK cell and T cell subsets was decreased in COVID-19 patients (Fig. 1a, b). Unbiased analysis of single chemokine receptors revealed a loss of CXCR2^+^, CXCR3^+^, CXCR6^+^, and CCR2^+^ cells despite an increase in the frequency of CCR5^+^ NK cells and T cells, respectively (Fig. 1c). When compared to influenza patients, we observed only minor and non-significant differences for chemokine receptor expression on NK cells in both diseases (Fig. 1d, e). Notably, a strong trend towards a loss of CXCR3^+^ NK cells in COVID-19 and influenza patients was observed (Fig. 1d, e). Lower frequencies of CXCR6^+^ NK cells were accompanied with an increase in CCR5^+^CD56^bright^CD16^-^ NK cells in influenza but not in COVID-19 patients (Fig. 1d, e). In contrast to NK cells, in particular CD8^+^ T cells were significantly affected both in COVID-19 and influenza patients (Fig. 1f, g). Similar to NK cells, CXCR3^+^ T cells were markedly reduced most, both in COVID-19 and influenza. Additionally, CXCR6^+^CD8^+^ T cells were reduced in influenza (Fig. 1f, g). Changes in chemokine receptor expression were observed both in naïve and non-naïve CD4^+^ and CD8^+^ T cells (Fig. S2a, b; gating strategy in Fig. S1a) and likewise when CD8^+^ TCM, TEM, and TEMRA cells were compared (Fig. S2c). Unbiased principal component analysis (PCA) revealed segregation between healthy controls and influenza patients for NK cells as well as between COVID-19 and influenza patients for T cells, respectively (Fig. 1h-j). The differences were largely driven by CXCR3 and CCR5 for NK cells and by CCR5 on CD4^+^ T cells as well as by CCR2, CXCR3, and CXCR6 on CD8^+^ T cells (Fig. 1i). Together, despite differences between NK cells and CD8^+^ T cells, both lymphocyte subsets displayed a similar pattern in terms of CXCR3 and CXCR6 expression in COVID-19 and influenza patients, with T cells being more strongly affected.

**Figure 1:**
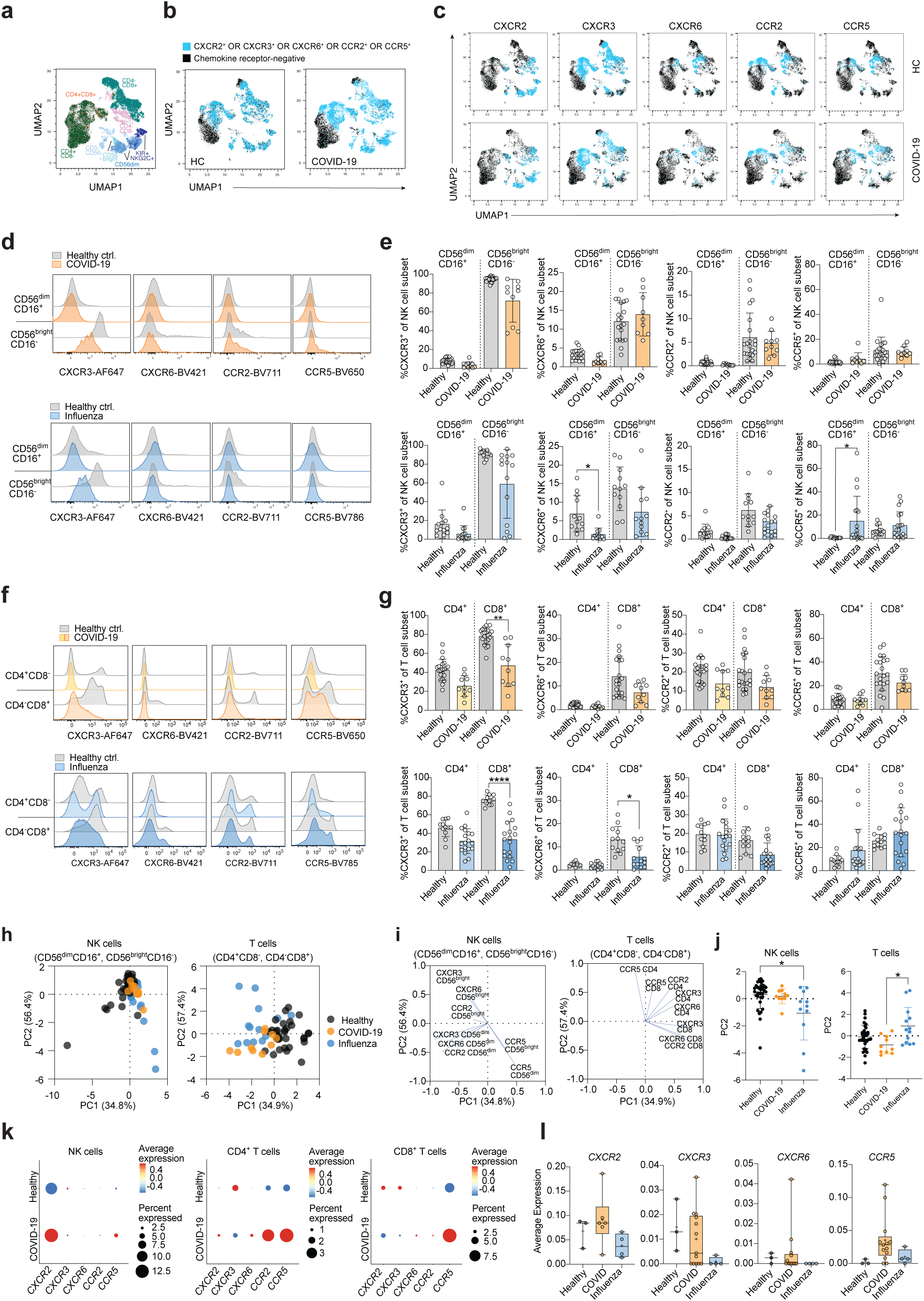
Altered composition of NK cells and T cells expressing different lung-homing receptors in peripheral blood in acute viral infection. **(a)** Uniform manifold approximation and projection (UMAP) analysis of NK and T cell subsets in healthy controls (HC) and COVID-19 patients. 15 healthy controls and 10 COVID-19 patients were included in the analysis with 3,000 cells of each donor. Concatanated HC data were downsampled to 30,000 events. **(b)** Chemokine receptor-positive cells (blue) in HC (left plot) or COVID-19 patients (right plot) were defined by applying a Boolean algorithm (CXCR2^+^ OR CXCR3^+^ OR CXCR6^+^ OR CCR2^+^ OR CCR5^+^) on data from (a). **(c)** Receptor-positive lymphocytes in HC (upper panel) and COVID-19 patients (lower panel) are indicated in blue, using the dataset from (a). **(d, f)** Representative overlays and **(e, g)** summary of data of chemokine receptor expression on **(d, e)** CD56^dim^CD16^+^ and CD56^bright^CD16^-^ blood NK cells in HC and COVID-19 patients (upper panel) or influenza patients (lower panel). (CXCR3: n = 14; CXCR6: n = 12; CCR2: n = 18; CCR5: n = 18. HC: n = 12), or on **(f, g)** CD4^+^CD8^-^ and CD4^-^CD8^+^ T cells in peripheral blood of HC and COVID-19 patients (n = 10) (upper panel) or influenza patients (lower panel; CXCR3: n = 18; CXCR6: n = 13; CCR2: n = 18; CCR5: n = 18. HC: n = 12). **(h)** PCA plots showing the distribution and segregation of healthy controls and COVID-19 and influenza patients, respectively based on expression of chemokine receptors (CXCR3, CXCR6, CCR2, CCR5) on CD56^bright^CD16^-^ and CD56^dim^CD16^+^ NK cells (left) and CD4^+^ and CD8^+^ T cells (right), respectively. Each dot represents one donor. **(i)** PCA plots showing corresponding trajectories of key receptors on NK and T cell subsets that influenced the group-defined segregation. **(j)** Dot plot showing the distribution of donors in PC2 for NK cells (left) and T cells (right). **(k)** Transcript expression of chemokine receptors in peripheral blood of HC (n = 3) and patients with moderate COVID-19 (n = 13) and influenza (n = 4) in NK cells (left), CD4^+^ T cells (middle), and CD8^+^ T cells (right). **(l)** Dot plots showing the average transcript expression of chemokine receptors on NK cells from healthy controls and COVID-19 and influenza patients. (k, l) Dataset derived from a publicly available scRNA-seq dataset (7). (e, g, j) Kruskal-Wallis rank-sum test with Dunn’s post hoc test for multiple comparisons. *p<0.05, **p<0.005, ****p<0.0001.

In order to compare the present phenotypic results, with those of other data collections, we analyzed two publicly available scRNA-seq datasets from peripheral blood from a different cohort of SARS-CoV-2-infected patients with moderate COVID-19 disease (6) (Fig. 1k) and from patients infected with either SARS-CoV-2 or influenza A virus (IAV) (7) (Fig. 1l), respectively. In order to allow a fair comparison of the RNA-seq data to the present dataset, we selectively analyzed data from patients with similar clinical characteristics. This analysis revealed differences of chemokine receptor expression between NK cells and T cells at the mRNA level in peripheral blood. While *CXCR2* was largely confined to NK cells, *CCR2* and *CCR5* dominated in T cells, both in healthy controls and in COVID-19 patients (Fig. 1k) (6). Lower average expression of *CXCR3* was found in all three lymphocyte subsets (NK cells, CD4^+^ T cells, CD8^+^ T cells) in COVID-19 patients as compared to healthy controls (Fig. 1k). However, in contrast to protein data, *CXCR6, CCR2*, and *CCR5* were strongly increased both in frequency and in mean intensity in T cells in COVID-19 patients as compared to healthy controls (Fig. 1k), suggesting post-transcriptional regulation of these chemokine receptors. Finally, direct comparative analysis of transcript expression of *CXCR2, CXCR3, CXCR6*, and *CCR5* revealed stronger loss of *CXCR2, CXCR3* and *CXCR6* in NK cells from influenza patients as compared to COVID-19 patients as well as a trend towards upregulation of *CCR5* (Fig. 1l) (7), in line with the similarly observed trend at the protein level (Fig. 1d, e) and suggesting a role for these chemokine receptors particularly in influenza infection. While CXCR3, CXCR6, CCR2, and CCR5 are predominantly expressed on CD56^bright^CD16^-^ NK cells in peripheral blood from healthy controls and patients with COVID-19 or influenza, respectively, CXCR2 is strongly expressed on CD56^dim^CD16^+^ blood NK cells (Fig. S2d, e). In COVID-19 patients, CXCR2^+^CD56^dim^CD16^+^ NK cells were mainly lost within the least differentiated NKG2A^+^CD57^-^ subset (Fig. S2f). On T cells, CXCR2 expression was overall low, without significant differences between healthy controls and COVID-19 patients (Fig. S2g, h).

Together, the present phenotypic analyses indicate that CXCR3 is a common lung-homing receptor for both NK cells and T cell subsets during acute infection with COVID-19 or influenza, despite major differences for *CXCR6, CCR2*, and *CCR5* between the two diseases at transcriptional level. Furthermore, CXCR6^+^ NK and T cells were strongly affected in influenza but not in COVID-19 patients, indicating potential differences in homing capacities in the two diseases.

### Activation profiles in chemokine-receptor positive NK cells differ in COVID-19 and influenza

We and others previously demonstrated an activated phenotype in peripheral blood NK cells and T cells in COVID-19 and influenza, respectively (4,9,10). Here, we aimed at identifying NK and T cell activation markers in relation to expression of lung-homing receptors as identified by boolean gating, combining cells expressing either CXCR3, CXCR6, CCR2, or CCR5 (Fig. S1b, c). Expression of activation markers such as CD69, CD38, Ki67, and NKG2D was elevated on NK cells in moderate COVID-19 patients (Fig. 2a, b), with strongest increases detected for CD69 and Ki67 (Fig. 2b, c). In detail, induction of CD69 expression was highest on CD56^bright^CD16^-^ NK cells co-expressing lung-homing receptors, while Ki67 induction was highest in corresponding CD56^dim^CD16^+^ NK cells, particularly in those co-expressing lung-homing receptors (Fig. 2c). Due to very low numbers of CD56^bright^CD16^-^ NK cells lacking any relevant chemokine receptor in healthy controls, no comparisons could be performed for this subset. In contrast to NK cells from COVID-19 patients, upregulation of CD69 was highest in CD56^dim^CD16^+^ NK cells in influenza patients, irrespective of co-expression of lung-homing receptors (Fig. 2d, e, f). Expression and upregulation of CD38 was similar in COVID-19 and influenza patients (Fig 2c, f). While these data indicate fundamental differences in activation patterns for NK cells in COVID-19 and influenza patients, other phenotypic characteristics remained stable between the two diseases (Fig. S3a, b), despite induction of granzymes and perforin observed in particular in chemokine receptor-positive CD56^bright^CD16^-^ NK cells (Fig. S3c-f). In the latter subset, upregulation of perforin expression was three times higher in influenza patients as compared to COVID-19 patients (Fig. S3e, f), indicating stronger activation of blood NK cells in influenza.

**Figure 2:**
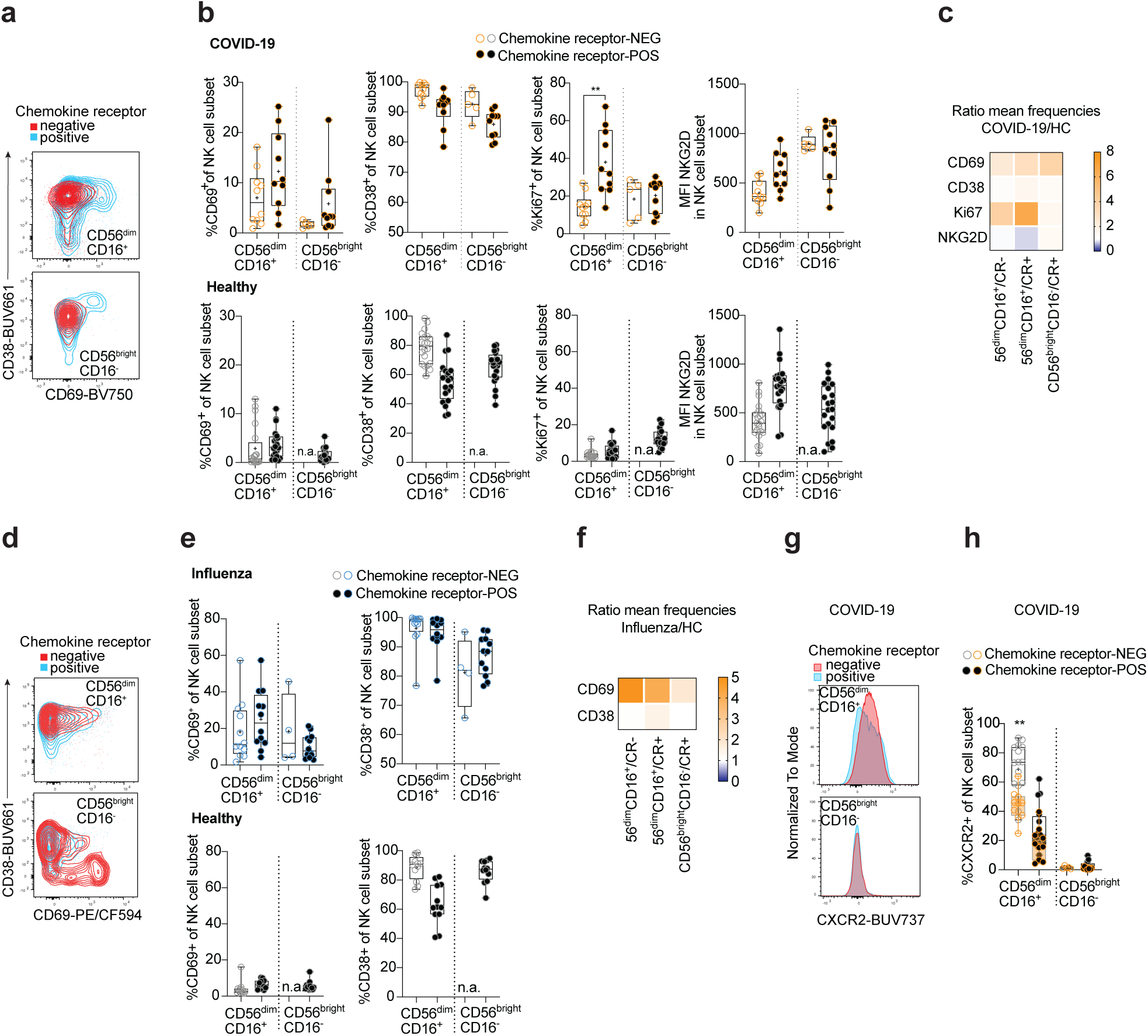
Lung-homing receptor-positive NK cells display an activated phenotype in peripheral blood of COVID-19 and influenza. Chemokine receptor-positive cells in COVID-19 and influenza patients were identified using the Boolean gate “CXCR3^+^ OR CXCR6^+^ OR CCR2^+^ OR CCR5^+^” (see representative gates in Figure S1b). Cells lacking all of these receptors were identified as chemokine receptor-negative. **(a)** Representative overlays and **(b)** summary of data showing CD69 and CVD38 on chemokine receptor-positive/negative CD56^dim^CD16^+^ and CD56^bright^CD16^-^ blood NK cells from COVID-19 patients (upper panel; n = 5-10) and healthy controls (lower panel; n = 20). **(c, f)** Heatmaps displaying the ratio of mean expression of **(c)** CD69, CD38, Ki67, and NKG2D between COVID-19 patients and healthy controls or **(f)** of CD69 and CD38 between influenza patients and healthy controls in CR^-^ and CR^+^ CD56^dim^CD16^+^ and CR^+^ CD56^bright^CD16-NK cells. Baseline value = 1 (white). **(d)** Representative overlays and **(e)** summary of data of CD69 and CD38 expression on CR^+^ and CR^-^ CD56^dim^CD16^+^ and CD56^bright^CD16^-^ blood NK cells from influenza patients (upper panel; n = 4-12) and healthy controls (lower panel; n = 12), respectively. **(g)** Representative overlays and **(h)** summary of data of CXCR2 expression on CR^-^ and CR^+^ CD56^dim^CD16^+^ and CD56^bright^CD16^-^ NK cells from COVID-19 patients (orange; n = 5) and healthy controls (grey; n = 16), respectively. Box and Whiskers, min to max, mean shown as ‘+’. Kruskal-Wallis rank-sum test with Dunn’s post hoc test for multiple comparisons. **p<0.005.

As indicated by transcript level, CXCR2 might have an additional role in lung-homing for NK cells. Since activation affected both CD56^dim^ and CD56^bright^ NK cells in COVID-19 patients, we next sought to determine changes in CXCR2^+^ expression (Fig. 2g, h). In this regard, expression of CXCR2 was higher on CD56^dim^CD16^+^ NK cells lacking other lung-homing receptors (Fig. 2g). The frequency of this CXCR2^+^CD56^dim^CD16^+^ NK cell subset was significantly reduced in COVID-19 patients as compared to healthy controls (Fig. 2h), indicating an alternative migration mechanism for CXCR2^+^CD56^dim^CD16^+^ NK cells to the lung in COVID-19 patients.

Together, our data show that the activation patterns differ for NK cell subsets in COVID-19 and influenza patients, respectively, with generally stronger activation of blood NK cells in influenza as compared to moderate COVID-19 patients. Furthermore, the data suggest that CXCR2 might act as an alternative lung-homing receptor for CD56^dim^CD16^+^ NK cells lacking other lung-homing receptors.

### Biased activation of T cells co-expressing lung-homing receptors in COVID-19 and influenza

Since NK cell subset activation in COVID-19 and influenza is associated with chemokine receptor expression, we next determined whether T cells expressing lung-homing receptors displayed an equivalent phenotypic bias towards stronger activation of chemokine receptor-positive T cells (Fig. 3). Similar to NK cells, a larger proportion of CD4^+^ and CD8^+^ T cells expressed CD69, both in COVID-19 (Fig. 3a, c) and influenza (Fig. 3b, c) compared to healthy controls. While in COVID-19 patients CD69 upregulation was biased towards CD8^+^ T cells co-expressing lung-homing receptors (Fig. 3a, c), upregulation was overall higher and more uniform between the T cell subsets in influenza (Fig. 3b, c). Furthermore, CD38 was strongly upregulated on CD8^+^ T cells in influenza but not COVID-19 patients (Fig. 3c). Finally, expression of Ki67 was largely confined to chemokine receptor-positive T cells, both in healthy controls and in COVID-19 patients (Fig. 3a). Upregulation of Ki67 was strongest in chemokine receptor-positive CD8^+^ T cells (Fig. 3c), which is in line with strongest upregulation in CD56^dim^CD16^+^ chemokine receptor-positive NK cells (Fig. 2c), indicating a particular activation of cytotoxic lymphocytes expressing lung-homing receptors in COVID-19 patients.

In comparison to NK cells where upregulation of granzymes and perforin was more uniform between COVID-19 and influenza patients (Fig. S3), differences were more distinct for T cells (Fig. 3d-f). As to be expected, expression of cytotoxic effector molecules was to a large extent contained to cytotoxic CD8^+^ T cells (Fig. 3d, f), although some expression was also observed in CD4^+^ T cells both in COVID-19 and influenza patients, respectively, with a particular bias towards chemokine receptor-negative CD4^+^ T cells (Fig. 3e, f). Expression of granzyme B and perforin was highest in chemokine receptor-negative CD8^+^ T cells in healthy controls as well as in COVID-19 and influenza patients (Fig. 3d, e). Importantly, however, lung-homing receptor-positive T cells displayed the highest increase compared to the same subset in healthy controls (Fig. 3f).

**Figure 3:**
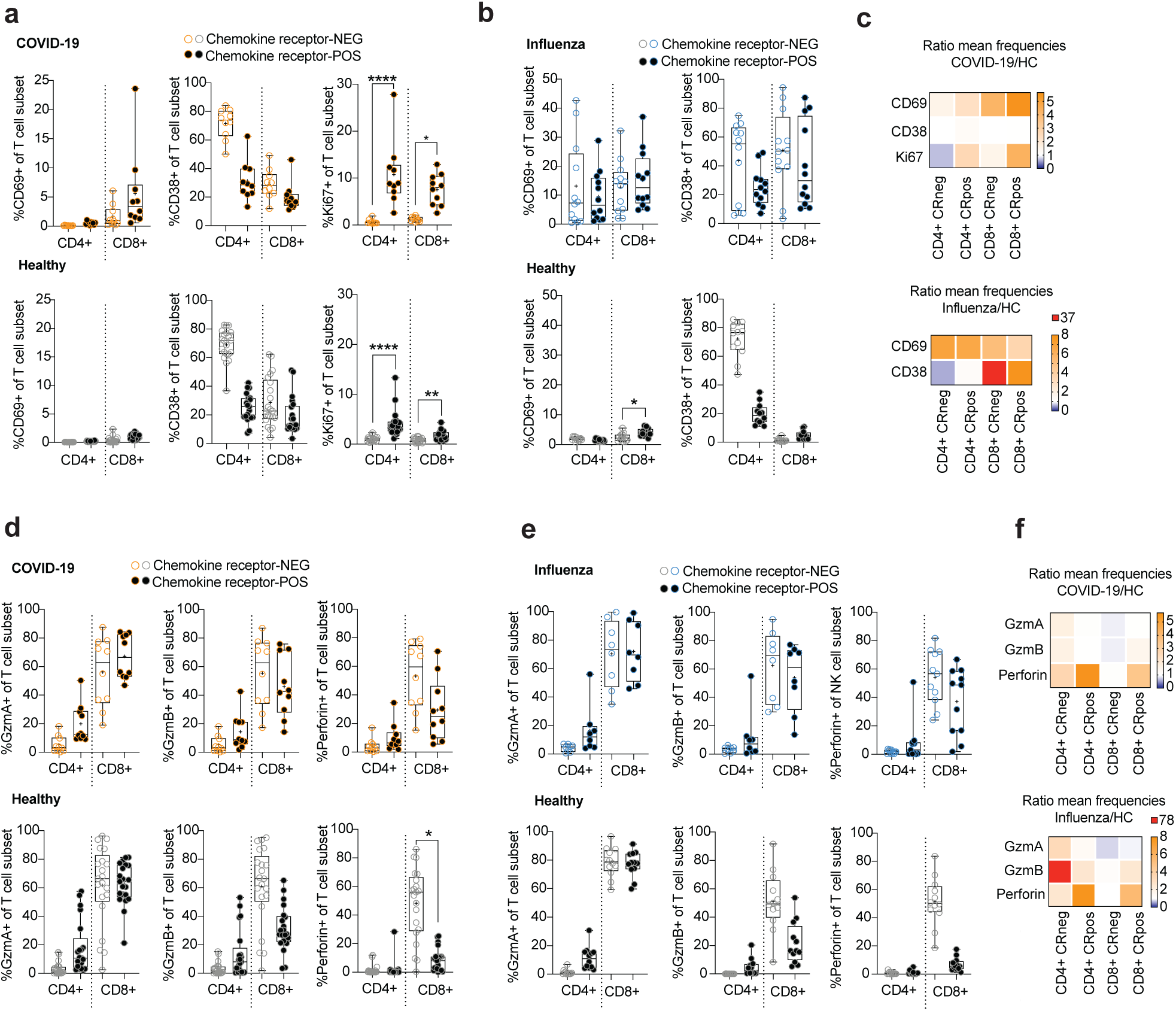
Circulating lung-homing receptor-positive T cells are phenotypically activated in COVID-19 and influenza. **(a, b)** Expression of **(a)** CD69, CD38, and Ki67 on CR^-^ and CR^+^ CD4^+^ and CD8^+^ T cells from COVID-19 patients (upper panel; n = 10) and healthy controls (lower panel; n = 21) or **(b)** of CD69, and CD38 on CR^-^ and CR^+^ CD4^+^ and CD8^+^ T cells from influenza patients (upper panel; n = 12) and healthy controls (lower panel; n = 12). **(c)** Heatmaps displaying the ratio of mean expression of CD69, CD38, and Ki67 between COVID-19 patients and healthy controls (top) or of CD69 and CD38 between influenza patients and healthy controls (bottom) in CR^-^ and CR^+^ CD4^+^ and CD8^+^ T cells. Baseline value = 1 (white). **(d, e)** Expression of effector molecules in CR^-^ and CR^+^ CD4^+^ and CD8^+^ T cells from **(d)** COVID-19 patients (upper panel; n = 10) and healthy controls (lower panel; n = 21), or **(e)** from influenza patients (upper panel; n = 8 (GzmA/B), n = 11 (perforin)) and healthy controls (lower panel; n = 12). **(f)** Heatmaps displaying the ratio of mean expression of effector molecules between COVID-19 patients (top) or influenza patients (bottom) and healthy controls in CR^-^ and CR^+^ CD4^+^ and CD8^+^ T cells. Baseline value = 1 (white). (a, b, d, e) Box and whiskers, min to max, mean is indicated as ‘+’. Kruskal-Wallis rank-sum test with Dunn’s post hoc test for multiple comparisons. *p<0.05, ****p<0.0001.

Together, these data indicate an overall stronger *in vivo* priming of T cells and NK cells in influenza patients as compared to moderate COVID-19 patients and also suggest a specific role for cytotoxic lymphocytes co-expressing lung-homing receptors.

### Distinct CXCR3- and CXCR6-mediated accumulation of phenotypically primed NK cells and T cells in BAL fluid of COVID-19 patients

Activation of cytotoxic NK cells and CD8^+^ T cells in COVID-19 and influenza patients was associated with high expression of effector molecules (Fig. S3, Fig. 3). Stratification of cells based on expression of chemokine receptors, granzyme A, granzyme B, and perforin revealed distinct co-expression patterns between CD56^bright^CD16^-^ and CD56^dim^CD16^+^ NK cells and CD8^+^ T cells as well as between COVID-19 and influenza (Fig. 4a). Shared between the different subsets and between COVID-19 and influenza was an increase of all effector molecules particularly on chemokine receptor-positive cells. Increased expression of effector molecules could also be confirmed at transcriptional level both for NK cells (Fig. 4b) and CD8^+^ T cells (Fig. 4c) in peripheral blood of COVID-19 patients as compared to healthy controls (6). Interestingly, levels of effector molecules gene transcripts were nearly identical between ventilated (severe) and non-ventilated (moderate) COVID-19 patients, both in NK cells (Fig. 4b) and CD8^+^ T cells (Fig. 4c), indicating that similar NK and T cell activation is similar in patients with moderate and severe disease.

**Figure 4:**
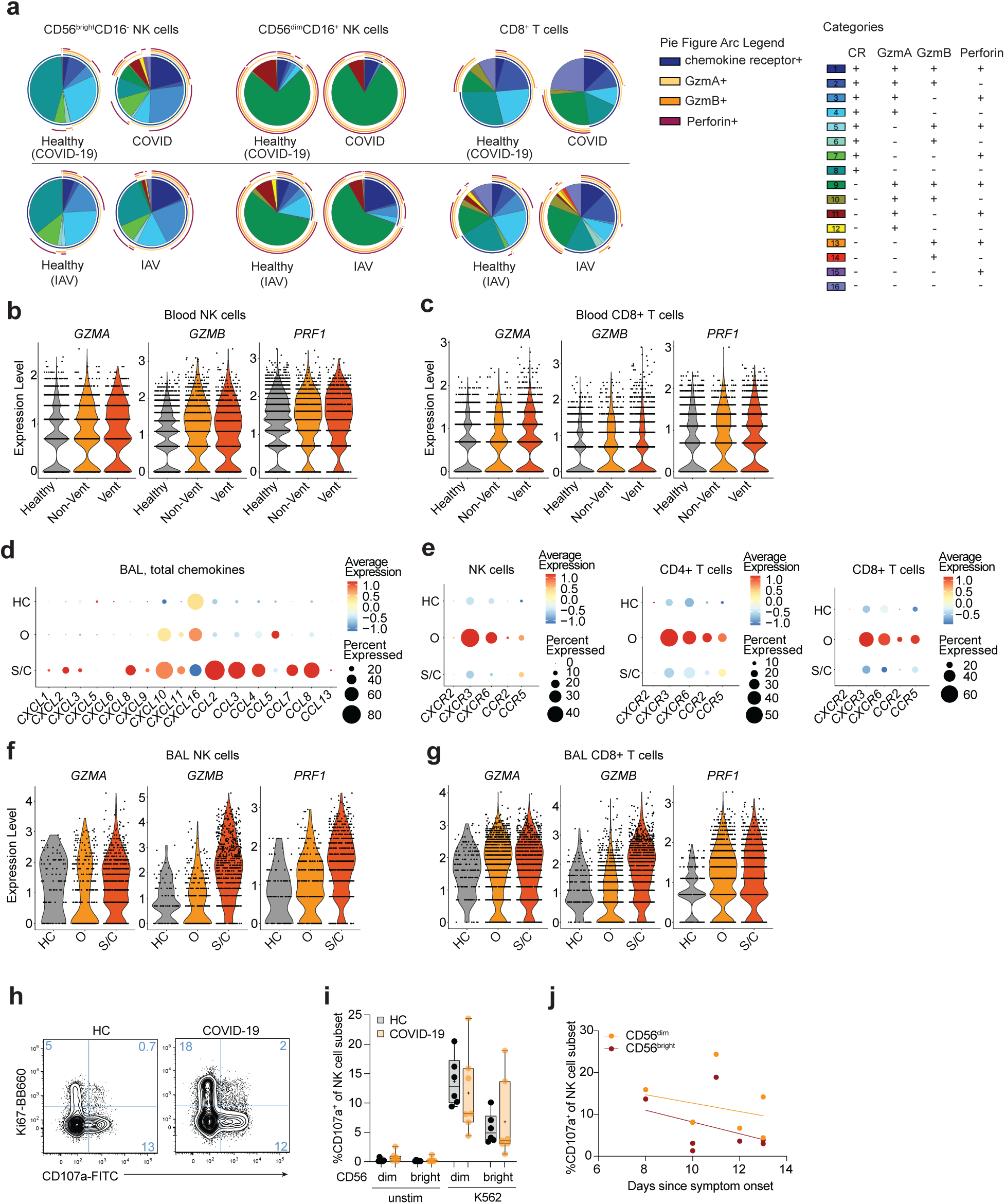
Upregulation of cytotoxic profile in circulating NK cells and CD8^+^ T cells in COVID-19. **(a)** SPICE analysis of CD56^bright^CD16^-^ NK cells (left), CD56^dim^CD16^+^ NK cells (middle), and CD8^+^ T cells (right) in COVID-19 patients (upper panel) and influenza patients (lower panel) and the respective healthy controls, displaying co-expression of effector molecules in CR^-^ and CR^+^ cells. n = 10/21 (COVID/healthy), n= 6/12 (influenza/healthy) **(b)** Violin plots of indicated genes encoding granzyme A, granzyme B, and perforin in NK cells and **(c)** CD8^+^ T cells from peripheral blood of healthy controls and non-ventilated and ventilated COVID-19 patients (6). **(d)** Dot plots of indicated genes in total BAL fluid cells, and **(e)** NK cells as well as CD4^+^ and CD8^+^ T cells from healthy controls and moderate and severe COVID-19 patients. **(f)** Violin plots of indicated genes in BAL fluid NK cells and **(g)** in CD8^+^ T cells in healthy controls and moderate and severe COVID-19 patients. **(h)** Representative dot plots showing CD107a and Ki67 expression in bulk NK cells from peripheral blood of a healthy control (HC) and a COVID-19 patient following stimulation with K562 cells. **(i)** Frequencies of CD107a expression in CD56^bright^ and CD56^dim^ NK cells of healthy controls (n = 6) or COVID-19 patients (n = 7) following 6 hour-incubation with or without K562 target cells. **(j)** Correlation analysis of CD107a expression frequencies in CD56^dim^ and CD56^bright^ NK cells (n = 7) versus time since symptom onset in COVID-19 patients. (d-g) RNAseq dataset derived from Liao *et al*. (8).

The loss of NK cells and T cells expressing lung-homing receptors in the peripheral blood of COVID-19 patients suggests migration of the respective cells to the infected lung tissue. In order to identify characteristics of NK cells and T cells in the lung, we next used a publicly available scRNAseq dataset from BAL fluid cells from COVID-19 patients (8) and analyzed expression of transcripts of relevant chemokines in total BAL cells (Fig. 4d) as well as chemokine receptor expression on NK cells and CD8^+^ T cells (Fig. 4e). High transcript levels of a large number of chemokines were found in patients with severe disease, while moderate patients were mainly distinguished from healthy patients by a significantly upregulated expression of *CXCL10* and *CCL5*, in addition to increased levels of *CXCL11* and *CXCL16* (Fig. 4d), highlighting the role for CXCR3, CXCR6, and CCR5 in moderate COVID-19. In line with these results, transcripts for *CXCR3, CXCR6*, and *CCR5* were highly enriched in NK cells as well as CD4^+^ and CD8^+^ T cells in COVID-19 patients with moderate disease (Fig. 4e). Although it is possible that some of the cells in BAL fluid are comprised of tissue-resident NK cells and memory T cells which express high levels of CXCR3 and CXCR6 at the transcriptional and protein levels (11,12), our data strongly suggest infiltration of NK cells and T cells from peripheral blood into the lung in COVID-19 patients. Since NK cells and CD8^+^ T cells from peripheral blood displayed upregulated levels of effector molecules, these cells might have important cytotoxic implications upon infiltration into the lung. Indeed, NK cells and CD8^+^ T cells in BAL fluid from COVID-19 patients displayed increased expression levels of *GZMA, GZMB*, and *PRF1* in patients with moderate and severe disease (Fig. 4e, f). We previously demonstrated that exposure to IAV-infected cells functionally primes human blood and lung NK cells *in vitro* towards increased target cell-responsiveness (9). However, despite phenotypic activation and upregulation of effector molecules both at RNA and protein levels, blood NK cells from patients with moderate COVID-19 showed no elevated responses to K562 target cells as compared to healthy controls (Fig. 4h, i). Instead, NK cell responses were slightly reduced with time after symptom onset (Fig. 4j). While our data suggest a low state of NK cell activation in peripheral blood of patients with moderate COVID-19, it remains to be determined whether NK cells migrating towards the lungs remain functional and contribute to lysis of virus-infected cells and possibly even to tissue pathology in patients with moderate or severe COVID-19.

Altogether, our data suggest distinct recruitment of CXCR3^+^ and CXCR6^+^ NK cells and CD8^+^ T cells to the lung in patients with moderate COVID-19 and influenza, indicating overlapping recruitment mechanisms in these two respiratory viral infections. Hence, a better understanding of lung-homing of innate and adaptive cytotoxic lymphocytes in COVID-19 and influenza patients might reveal universal concepts of disease progression in these two, and possibly other, respiratory viral infections.

## Discussion

The COVID-19 pandemic has raised the awareness about the need for a better understanding of the course of respiratory viral infections. So far, only a few studies have compared immune responses in COVID-19 and other respiratory viral infections side by side (13) (14,15). Here, we compared COVID-19 with influenza. Both viruses are to different degrees transmitted by contact, droplets and fomites. They cause respiratory disease with similar disease presentation ranging from asymptomatic or mild to severe disease and death. Here, we demonstrate that NK cells and T cells, in particular CD8^+^ T cells, largely overlap in their chemokine receptor expression profile in the blood of COVID-19 and influenza, indicating similar lung-homing mechanisms for cytotoxic lymphocytes in both infections. We identified a stronger loss of CXCR3^+^ and CXCR6^+^ CD8^+^ T cells, overall stronger activation of NK cells and T cells, as well as reduced transcript level expression of lung-homing receptors in the blood of influenza patients as compared to COVID-19 patients with moderate disease, in line with the recent observation of a lower inflammatory profile in COVID-19 patients as compared to influenza patients (14). The activated profile of NK cells and CD8^+^ T cells, including elevated expression of CD69 and Ki67, and induction of perforin, was biased towards subsets co-expressing one or more of the lung-homing receptors CXCR2, CXCR3, CXCR6, CCR2, or CCR5. Levels of corresponding leukocyte-recruiting chemokines such as IL-8, CCL5, CXCL1, CXCL2, CXCL5, and CXCL10 (IP-10), are elevated in the BAL fluid of patients infected with SARS-CoV-2 (16,17). Monocytes producing ligands for CXCR3 have been shown to be expanded in the lungs of COVID-19 patients (8). Data regarding chemokine expression in the lung in influenza are limited to patients with fatal outcome (2). However, in particular CXCL10 has been shown to be upregulated in serum of influenza patients (18,19) and in SARS-CoV-1 patients with ARDS (20), in *in vitro* influenza virus-infected human macrophages (21), in human lung tissue explants infected with SARS-CoV-2 (16), and in the lungs of mice infected with influenza virus (3,22,23), suggesting a predominant role for the CXCR3:CXCL10 axis for lung-homing upon respiratory viral infection and, interestingly, also in non-viral lung tissue injury (24). Elevated CXCL10 levels in BAL was associated with longer duration of mechanical ventilation in COVID-19 patients (17). In mice, CXCR3-deficiency rescued CCR5-deficient mice from IAV-induced mortality (25). Other murine IAV infection models demonstrated a role for CXCR3 and CCR5 for NK cell lung-homing and showed NK cell accumulation in the lung was not due to proliferation or apoptosis (23). For CD8^+^ T cells, murine models demonstrated that virus-specific T cells express CXCR3 and migrate to CXCR3 ligands *in vitro* (26). Furthermore, CCR5 is required for recruitment of memory CD8^+^ T cells to IAV-infected epithelium and is rapidly upregulated on the surface of memory CD8^+^ T cells upon viral challenge (26). Interestingly, these mouse models also revealed that CCR5 is required for circulating CD8^+^ memory T cells to migrate to respiratory airways but not lung parenchyma during virus challenge (26), indicating potential distinct migration patterns depending on chemokine receptor expression. In addition to CCR5 and CXCR3, CXCR6 has been suggested to be of importance for recruitment of resident memory T cells to the airways both in mice (12) and in moderate COVID-19 (8).

The relevance of the CXCR3:CXCL10 axis for lung tissue-homing of cytotoxic immune cells such as NK cells and CD8^+^ T cells might be of interest for future approaches of intervention. Antibody-mediated targeting of CXCL10 improved survival of H1N1-infected mice (27), revealing a novel approach for immunotherapy in patients with severe respiratory viral infections. A thorough review recently summarized relevant known factors and cells involved in lung-homing during infection with SARS-CoV-2 and influenza, emphasizing the role of circulating NK cells not only in terms of their cytotoxic potential but also in potentially facilitating the recruitment of other cell types such as neutrophils (28). The potential immunoregulatory roles of NK cells in the human lung in health and disease however remains to be studied further.

Despite the parallels between SARS-CoV-2 and influenza virus infection, both diseases differ at an immunological level including that SARS-CoV-2 does not infect NK cells or other mononuclear blood cells due to the lack of ACE2 expression (29), while influenza virus can infect NK cells (30). In contrast to influenza virus, SARS-CoV-2 can spread to other organs if not cleared efficiently from the respiratory tract (31), and the viruses induce different antiviral responses in lung epithelial cells (32).

Here, we demonstrate common lung-homing potential of circulating NK cells and CD8^+^ T cells in SARS-CoV-2 and influenza patients. A shortcoming of our study is a low number of patients, impeding a detailed stratification by clinical or other parameters. However, the present cohort was limited to clinically well-defined patients with moderate disease. Future studies will assess how acute respiratory viral infection with SARS-CoV-2, influenza viruses, and other viruses affects the landscape of activated NK cells and T cells in the lung.

Together, our results strongly implicate an importance of CXCR3 as a lung-homing receptor in respiratory viral infections such as SARS-CoV-2 and influenza virus in humans. The results also reveal a role for other receptors such as CXCR6, CCR5 on CD56^bright^CD16^-^ NK cells, as well as CXCR2 on CD56^dim^CD16^+^ NK cells, as potential alternative receptors of importance. A better understanding of how these chemokine receptors are affecting disease progression might help to develop future immunotherapeutic interventions in patients that developed disease in current or future epidemics or pandemics with respiratory viral infections.

## Material and Methods

### Patients and processing of peripheral blood

We enrolled a total of 10 hospitalized patients (four females and six males; age range 24-70; average age 55.3) who were diagnosed with COVID-19 by RT-qPCR for SARS-CoV-2 in respiratory samples. Patients were sampled on average 11 days (range 6-16) after symptom onset. All of the COVID-19 patients were considered ‘moderate’ based on the guidelines for diagnosis and treatment of COVID-19 (Version 7) released by the National Health Commission and State Administration of Traditional Chinese Medicine (33). Furthermore, we enrolled 18 patients who tested positive for IAV (n=12) or IBV (n=6) (nine females and nine males; age range 21-84; median age 45) by RT-qPCR who were recruited during the four months immediately preceding the outbreak of COVID-19 in the Stockholm region. Nine of the 18 influenza patients were hospitalized.

None of the COVID-19 patients and only one of the influenza patients received immunosuppressive treatment. Diagnostics for all patients were performed at the diagnostic laboratory at the Karolinska University Hospital, Stockholm, Sweden. Patients were sampled on average 5 days (range 1-11) after symptom onset. Mononuclear cells from peripheral blood were isolated by density gradient centrifugation (Lymphoprep). For each of the two separate cohorts, blood was collected from healthy blood donors and processed in parallel with patient samples.

The study was approved by the Regional Ethical Review Board in Stockholm, Sweden, and by the Swedish Ethical Review Authority. All donors provided informed written consent prior to blood sampling.

### Transcriptome analysis

Preprocessed and annotated scRNA-seq datasets of cells from peripheral blood and BAL fluid from healthy controls, COVID patients, and influenza patients were obtained in RDS formats from published data (6-8). The data was read and analyzed using Seurat (v3.2.2). Quality filtering and cell and cluster annotations were retained from the respective publications. The datasets were scaled and transformed using the SCTransform function with mitochondrial gene expression regressed out, this was repeated after excluding populations and groups. Differentially expressed genes present in at least 25% of the cells were determined in a pairwise manner comparing patient groups using the sctransform outcome and the FindMarkers function implemented in Seurat. NK cells and T cell subsets were selected separately and analyzed based on patient group for transcript expression of chemokine receptors and effector molecules. CD4^+^ T cells and CD8^+^ T cells were identified from the BAL T cell subset based on expression of CD4 and CD8. NK cells and T cell populations were further studied in only moderate and non-ventilated COVID patients in the datasets from peripheral blood and BAL fluid, respectively.

### Flow cytometry

Antibodies and clones used for phenotyping are listed in supplementary table 1. Secondary staining was performed with streptavidin BUV630 (BD Biosciences) and Live/Dead Aqua (Invitrogen). After surface staining, peripheral blood mononuclear cells (PBMC) were fixed and permeabilized using FoxP3/Transcription Factor staining kit (eBioscience).

Samples were analyzed on a BD LSR Fortessa equipped with four lasers (BD Biosciences), and data were analyzed using FlowJo version 9.5.2 and version 10.7.1 (Tree Star Inc). UMAPs were constructed in FlowJo 10.7.1 using the UMAP plugin. For calculation of the UMAP, the following parameters were included: CXCR3, CCR7, GzmA, GzmB, Perforin, FcεR1g, NKG2C, Ki67, CD8, CD16, CD56, CD38, CXCR2, CD3, CXCR6, CD4, CD57, CCR5, CCR2, CD69, CD45RA, NKG2A, NKG2D, pan-KIR, NKp80.

### Degranulation assay

Cells were co-cultured in R10 medium alone or in presence of K562 cells for 6 hours in the presence of anti-human CD107a (H4A3, FITC, BD Biosciences). Golgi plug and Golgi stop (BD Biosciences) were added for the last 5 hours of the assay. The PBMC were subsequently stained as described above.

### Principal component analysis

Analyses were performed using GraphPad Prism 9 (GraphPad Software). PCs with eigenvalues greater than 1.0 (“Kaiser rule”) were used for component selection.

### Statistical analyses

GraphPad Prism 8 and 9 (GraphPad Software) was used for statistical analyses. The statistical method used is indicated in each figure legend.

## Author contributions

Conceptualization/study design: D.B., M.B., S.G.R., A.S.S., N.M.; Investigation: D.B., I.R., R.V., S.F.J., S.V., J.M., N.M.; Resources: H.A., H.G., S.F.J., S.V., S.G.R., A.S.S.; Writing – original draft: D.B., N.M.; Writing – review and editing: D.B, I.R., R.V., H.A., H.G., S.F.J., S.V., M.B., H.G.L., J.M., S.G.M., A.S.S., N.M.

## Acknowledgments

We thank the volunteers who participated in the study and A. v. Kries for assistance in the laboratory.

