## Supplemental Figure 1 for "Distinct lung-homing receptor expression and activation profiles on NK cell and T cell subsets in COVID-19 and influenza"

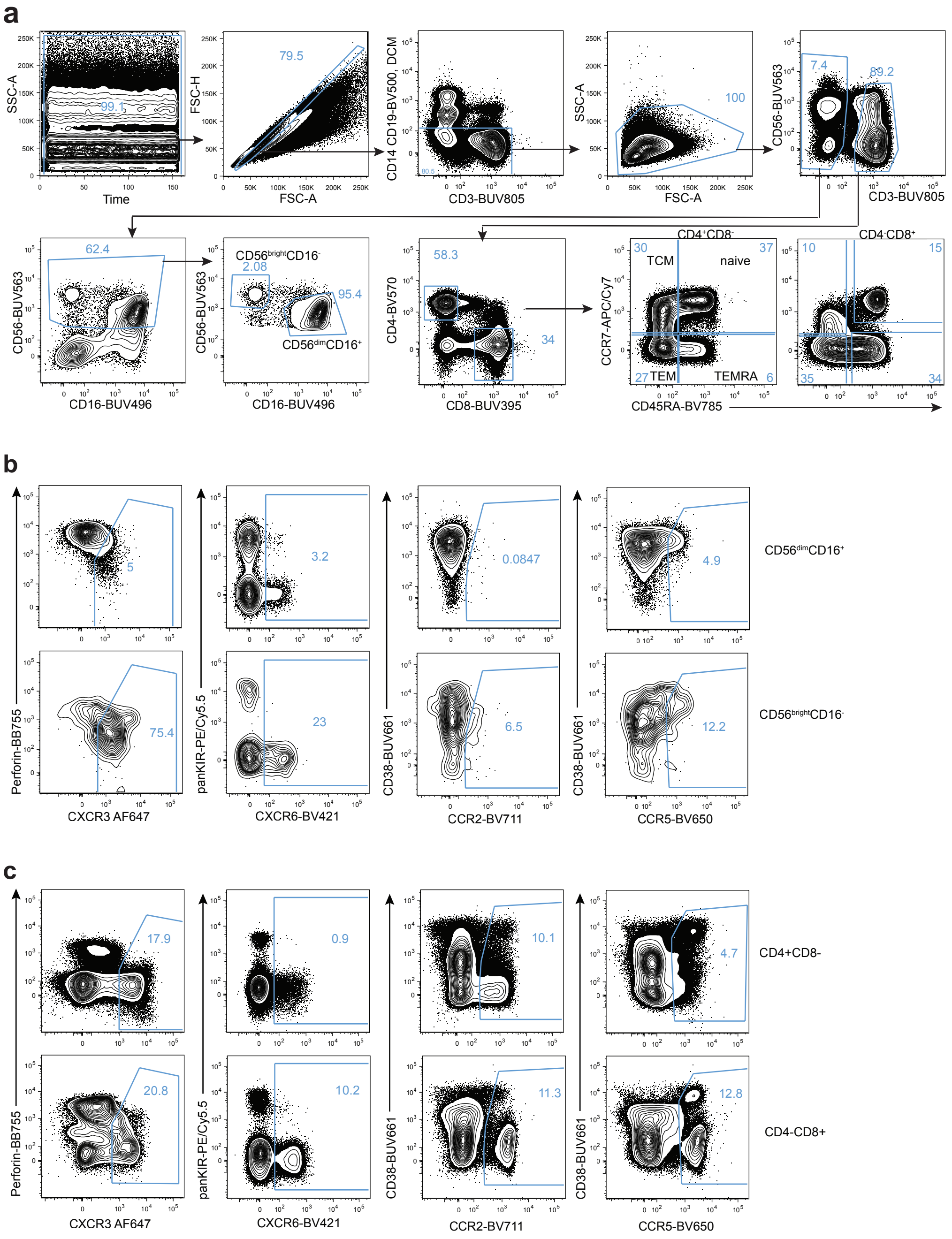


**Supplementary Figure 1: Gating strategies. (a)** Gating strategy for identification of NK cell and T cell subsets in peripheral blood of COVID-19 and influenza patients, respectively. **(b)** Representative dot plots showing gates for CCR2, CCR5, CXCR3, and CXCR6 in CD56$^{dim}$CD16$^{+}$ and CD56$^{bright}$CD16$^{-}$ NK cells or **(c)** CD4$^{+}$ and CD8$^{+}$ T cells, respectively, for the identification of chemokine receptor-positive cells in the respective subset. Chemokine receptor-positive cells were defined by a boolean algorithm ("CXCR3$^{+}$ OR CXCR6$^{+}$ OR CCR2$^{+}$ OR CCR5$^{+}$").
