## Supplemental Figure 2 for "Distinct lung-homing receptor expression and activation profiles on NK cell and T cell subsets in COVID-19 and influenza"

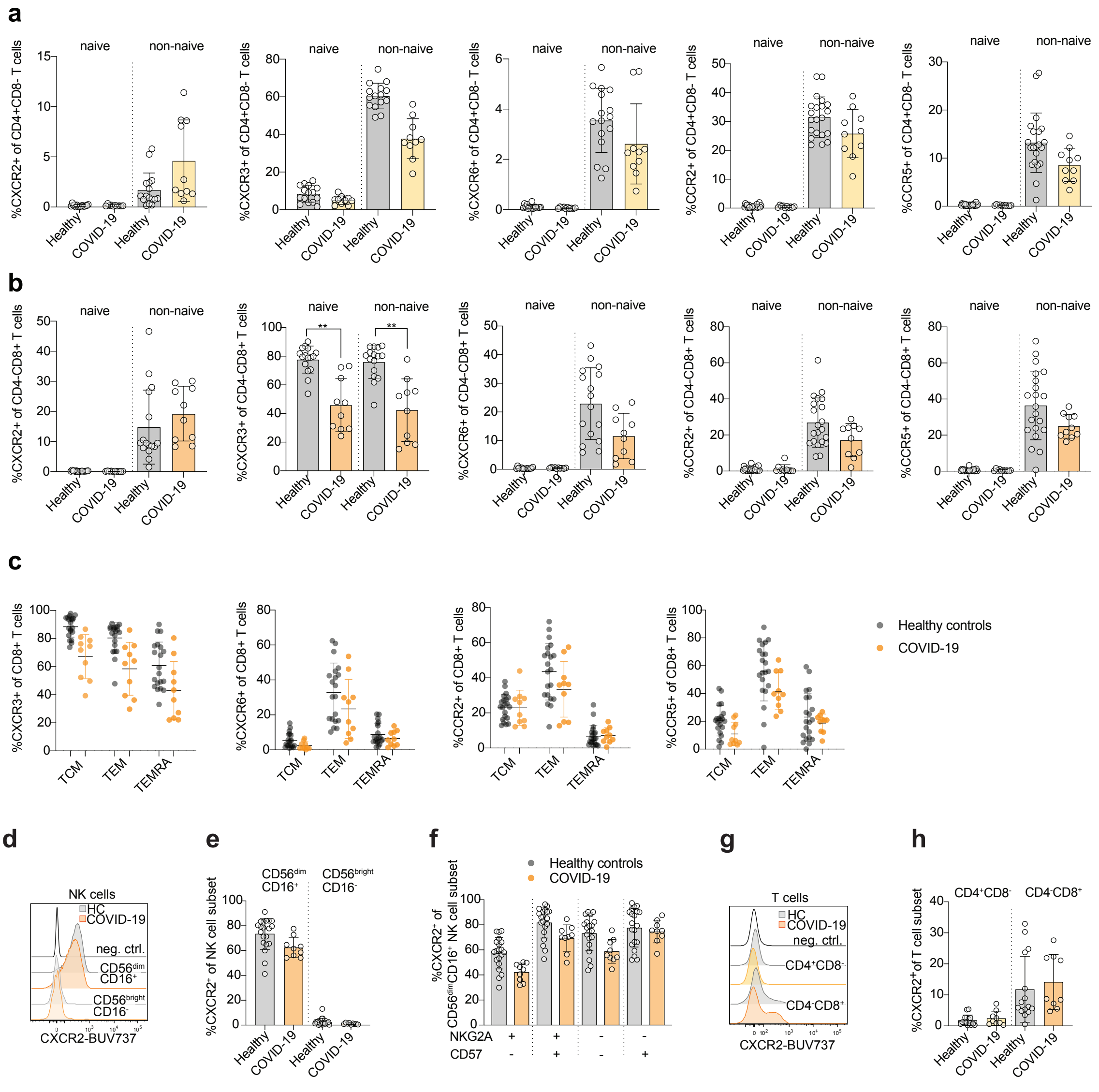


**Supplementary Fig. 2: Chemokine receptor expression on NK cell and T cell subsets in COVID-19.** **(a)** Expression of chemokine receptors on naive and non-naive $CD4^+$ and **(b)** $CD8^+$ T cell subsets. Kruskal-Wallis test. **p<0.001. **(c)** Frequencies of expression of chemokine receptors on non-naive $CD8^+$ T cell subsets. Healthy controls n = 21; COVID-19 n = 10. **(d)** Representative overlay and **(e)** summary of data of CXCR2 expression on $CD56^{dim}CD16^+$ and $CD56^{bright}CD16^-$ NK cells in healthy control (HC) and COVID-19 patients. **(f)** Summary of data of CXCR2 expression at distinct differentiation stages of $CD56^{dim}CD16^+$ NK cells in healthy controls (n = 20) and COVID-19 patients (n=9). **(g)** Representative overlay and **(h)** summary of data of CXCR2 expression on $CD4^+CD8^-$ and $CD4^-CD8^+$ T cells in HC (n = 15) and COVID-19 patients (n = 9).
