## Supplemental Figure 3 for "Distinct lung-homing receptor expression and activation profiles on NK cell and T cell subsets in COVID-19 and influenza"

### Supplementary Fig. 3

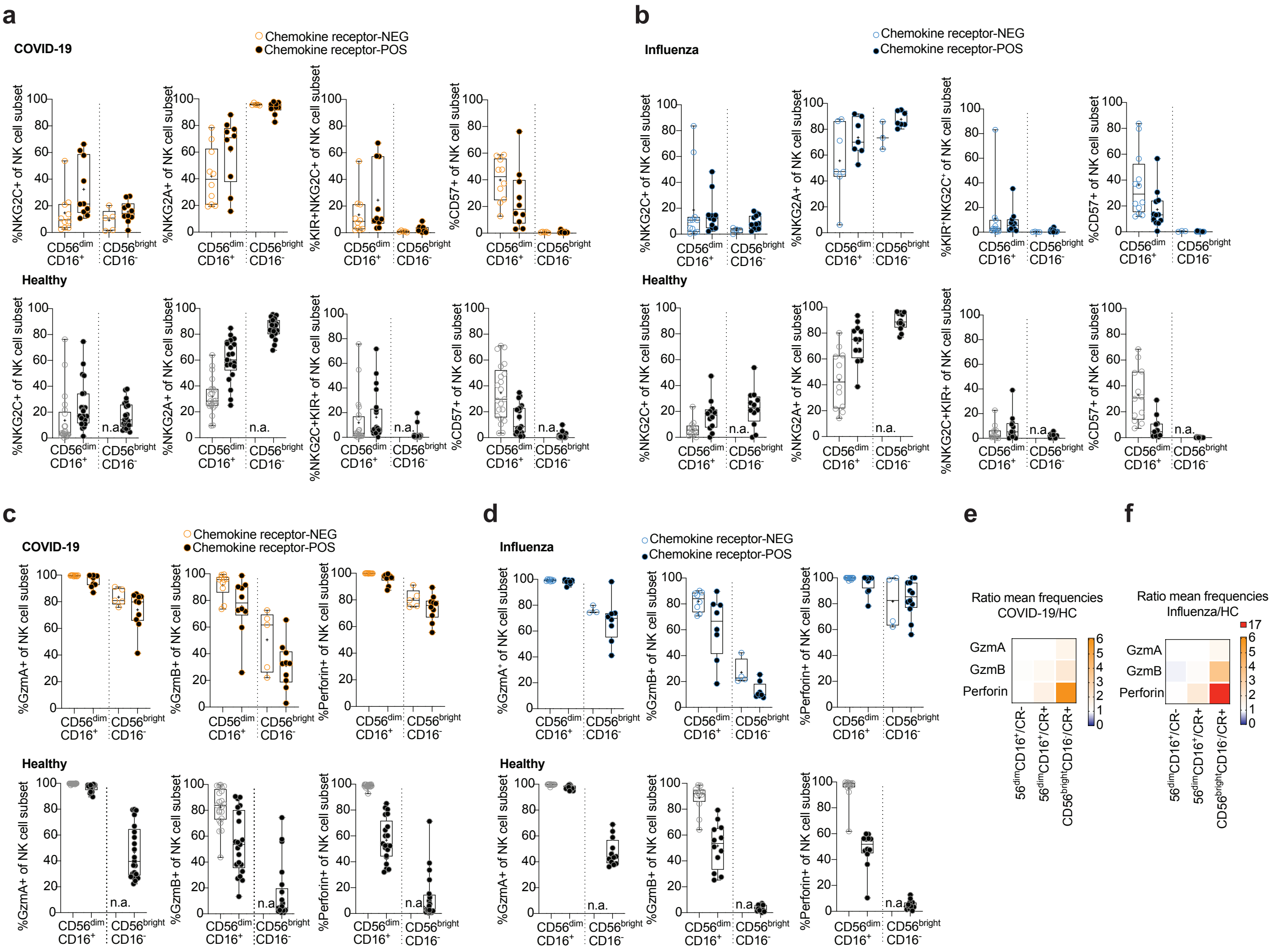


**Supplementary Fig. 3: Assessment of phenotypic alterations on NK cell subsets in COVID-19 patients and influenza patients.** **(a, b)** Frequencies of surface marker positive CD56$^{dim}$CD16$^{+}$ or CD56$^{bright}$CD16$^{-}$ NK cells in **(a)** COVID-19 patients (upper panel), **(b)** influenza patients (upper panel), and healthy controls (lower panels). **(c, d)** Frequencies of CD56$^{dim}$CD16$^{+}$ and CD56$^{bright}$CD16$^{-}$ NK cells co-expressing effector molecules in **(c)** COVID-19 patients (upper panel), **(d)** influenza patients (upper panel), and healthy controls (lower panels). a-d: Box and whiskers, min to max, mean is indicated as '+'. **(e, f)** Heatmaps displaying the ratio of mean frequencies of effector molecule expression in chemokine receptor (CR) negative and positive CD56$^{dim}$CD16$^{+}$ and CD56$^{bright}$CD16$^{-}$ NK cells, respectively, between **(e)** COVID-19 patients or **(f)** influenza patients and healthy controls (HC).
